# On pathogenicity with varying microbial virulence and host resistance

**DOI:** 10.1101/2024.10.02.616271

**Authors:** Anjali Gupta, Robert L. Unckless

## Abstract

The pathogenicity of a microbe is difficult to define in a comparative context: across microbes with varying virulence or across host genotypes with varying susceptibilities. Several different statistical approaches have been employed to investigate pathogenicity and susceptibility. Measures like proportional mortality or morbidity at a given time are attractive due to their simplicity but represent a single snapshot. Survival curve approaches, such as the Cox proportional hazards model and risk scores provide a more nuanced picture of the course of infection. More recently, Casadevall introduced the concept of pathogenic potential, a composite measure encompassing both host susceptibility and pathogen virulence, which focuses on the pathogenicity of a single pathogenic microbe, and later expanded to include additional nuances. Using *Drosophila melanogaster*, we conducted infection experiments with diverse species of *Providencia* bacteria that naturally vary in virulence. We also used several infectious doses to tune infections. We employed different host genotypes that vary in susceptibility to *Providencia* infection. Our analysis incorporates factors such as host genotype, pathogen type, inoculum load, symptomatic fraction, and mortality to compare host- and pathogen-based measures of pathogenicity. We discuss the advantages and limitations of each method, providing insights into their applicability in deciphering the intricacies of host-pathogen interactions and guiding the choice of analytical approaches in infectious disease research.

## Introduction

Virulence is the relative capacity of a microbe to cause damage in a host (Casadevall & Pirofski, 1999). It is intricately linked to host susceptibility, and its measurement is context-dependent, varying across host systems or for interspecific microbial comparisons (Casadevall & Pirofski, 2001). But virulence lacks an absolute measure, emphasizing the need for a relative assessment of a microbe’s impact on a host (Casadevall & Pirofski, 1999). Hosts may display symptoms when infected with a specific microbial threshold, exhibiting observable disruptions in normal physiological functions, including mortality in severe cases. This relationship is further complicated by the fact that virulence is not a static characteristic but can evolve in response to changes in host resistance, environmental conditions, and microbial interactions (Ewald, 1994; Read, 1994). Host resistance refers to the ability to limit pathogen burden, while tolerance describes the capacity to minimize damage caused by infection (Råberg et al., 2009). Understanding these distinctions is essential to interpreting the genetic and evolutionary underpinnings of infection outcomes. Pathogen virulence is similarly modulated by evolutionary trade-offs between transmission and host survival. For instance, highly virulent strains may dominate in environments where transmission opportunities are frequent and host mortality does not significantly impede pathogen spread (Frank, 1996). Conversely, in environments where hosts are scarce or transmission is limited, less virulent strains that prolong host survival may be favored (Ebert & Bull, 2003). These trade-offs illustrate the intricate evolutionary dynamics governing host-pathogen interactions and the maintenance of genetic variation within both host and pathogen populations. Additionally, the expression of virulence factors can be modulated by the host immune response, with some pathogens evolving mechanisms to evade or suppress host defenses, thereby enhancing their pathogenic potential (Casadevall & Pirofski, 2003; Mims et al., 2001). Understanding these dynamics is crucial for developing effective disease management strategies and predicting the emergence of more virulent or drug-resistant strains (Anderson & May, 1982; Antia et al., 2003).

Moreover, the interplay between microbial virulence and host resistance underscores the importance of considering both pathogen and host genetics in the study of infectious diseases. Genetic variations in hosts can influence susceptibility to infections, leading to diverse clinical outcomes even within the same species (Hill, 2001; Schmid-Hempel & Ebert, 2003). Similarly, microbial genetic diversity can result in different virulence profiles, impacting the severity and spread of infections (Mackinnon & Read, 2004; Woolhouse et al., 2002). Therefore, a comprehensive understanding of virulence requires an integrated approach that examines both microbial and host factors, as well as their interactions within specific ecological and evolutionary contexts (Casadevall & Pirofski, 2001; Galvani, 2003).

Infections often progress most critically during early time points, when host responses are most active, and pathogen replication is rapid. The early infection period is crucial in determining the trajectory of disease outcomes, as it captures the dynamic interplay between host defenses and pathogen strategies (Duneau et al., 2017). Metrics that focus solely on survival or endpoint measures may miss these critical early events, underscoring the need for more nuanced approaches to study host-pathogen dynamics. In many studies, the quantification of host susceptibility and pathogen virulence relies on host mortality rates, often using the Cox proportional hazards or related models (Cox, 1972; Klein & Moeschberger, 2003). This model offers a robust framework for survival analysis, combining a non-parametric baseline hazard with a parametric model for covariates proportional to the baseline hazard (Therneau et al., 2000). Risk scores, crucial in survival analysis, are calculated based on the Cox model (Hosmer Jr et al., 2008). While risk scores have been employed to assess risk factors, they do not encompass a quantitative measure of pathogen virulence and solely rely on host mortality (Bradburn et al., 2003).

Simple statistics such as the proportion of hosts alive at different time points and the time to death (or survival time) are commonly used to summarize survival data. The proportion alive provides a straightforward measure of survival at specific times but lacks the nuance to account for varying follow-up times and censoring (Aalen et al., 2008; Cutler & Ederer, 1958). Time to death offers more detailed information but is limited by the need for complete survival data, may not accurately reflect the impact of covariates (Altman, 1990) and is inappropriate for pathogens where hosts can make a full recovery. The Kaplan-Meier estimator is another widely used method in survival analysis that provides an estimate of the survival function from lifetime data (Kaplan & Meier, 1958). This non-parametric statistic handles censored data effectively and allows for the estimation of survival probabilities at various time points. However, it does not account for the effects of multiple covariates simultaneously, limiting its utility in multifactorial analyses (Collett, 2023; Pocock et al., 2002). To address these limitations, the Cox proportional hazards model offers a more comprehensive approach by incorporating multiple covariates into the survival analysis, thus providing insights into the factors that influence host mortality rates (Cox, 1972). However, while this model is powerful for identifying risk factors, it does not directly quantify pathogen virulence, necessitating additional methods to capture the full complexity of host-pathogen interactions (Hosmer Jr et al., 2008).

Casadevall, 2017 introduced the concept of the pathogenic potential of a (single) microbe, highlighting the variable nature of host damage during host-microbe interactions. Clinical symptoms manifest when host damage disrupts homeostasis, leading to death when damage surpasses the point of repair and return to homeostasis becomes impossible. Pathogenic potential is conceptualized as a composite of host susceptibility and pathogen virulence, expressed as 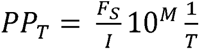, where PP represents pathogenic potential, F denotes the fraction symptomatic (host susceptibility factor), / is the infective dose (or pathogen burden), M is the mortality fraction and *T* represents time to disease manifestation (Casadevall, 2017). In this model, the fraction symptomatic host (F_S_) is a measure of host susceptibility, referring to the inherent or acquired traits of the host that affect its vulnerability to infection and disease progression. These traits can include genetic factors, immune status, and existing health conditions that influence how severely the host is affected by the pathogen (Casadevall, 2017). The infective dose (/) is a measure of pathogen virulence, pertaining to the ability of the pathogen to cause damage to the host, which is determined by its mechanisms of infection, replication rate, and production of virulence factors that harm the host (Casadevall, 2017). The mortality fraction (M) is a measure of the lethality of the infection, indicating the proportion of infected hosts that succumb to the disease (Casadevall, 2017). The time to disease (T) captures the duration from infection to the onset of clinical symptoms, adding a temporal dimension to the pathogenic potential and reflecting the speed of disease progression (Casadevall, 2017).

To fully understand the dynamics of host-pathogen interactions, is it crucial to distinguish between host-based and pathogen-based measures of pathogenicity? Or, do these metrics ultimately capture similar aspects of disease dynamics? Host-based measures focus on the response of the host organism to the pathogen, including metrics such as survival rates, risk scores, and the proportion of symptomatic individuals (Schmid-Hempel, 2009). These measures provide insight into how the host’s health and survival are affected by infection, often using tools like Kaplan-Meier survival curves and Cox proportional hazards models to analyze mortality data (Cox, 1972; Kaplan & Meier, 1958). But do they truly reflect the virulence of the pathogen itself, or are they more indicative of the host’s capacity to withstand infection (Kleinbaum & Klein, 1996)? In contrast, pathogen-based measures aim to quantify the inherent virulence of the pathogen, independent of the host’s response. This includes assessing the pathogen’s ability to cause disease at various doses and its potential to induce host mortality (Mideo et al., 2008; Read, 1994). Are these distinctions meaningful, particularly in cases of high-dose infections where it becomes difficult to attribute mortality to a specific pathogen due to overwhelming pathogen loads (Frank & Schmid Hempel, 2008)? By exploring these measures, can we better understand the relationship between host resilience and pathogen virulence, or are these categories less distinct than they initially appear?

Here, we set out to investigate the informativeness of quantitative measures of pathogenicity, including simple metrics of survival, cox survival analysis, and pathogen potential and its related measures. We conducted infections of *Drosophila melanogaster* using *Providencia* species as a pathogen due to their varied levels of virulence and interaction with the host immune system.

The *Providencia* genus offers a compelling system for investigating these dynamics. These bacteria exhibit significant genetic diversity in virulence, with some species causing highly lethal infections, while others result in moderate or minimal host damage (Galac & Lazzaro, 2011). *P. sneebia* and *P. alcalifaciens* are highly pathogenic to *D. melanogaster*, with even low doses leading to lethal infections (Galac & Lazzaro, 2011). In contrast, *P. burhodogranariea* Strain D exhibits lower virulence, making it a suitable model for studying infections that result in moderate survival rates (Galac & Lazzaro, 2011). This variation across species provides a powerful framework to examine the genetic and evolutionary mechanisms underlying differences in pathogen virulence and host susceptibility. *Providencia rettgeri* causes differential survival outcomes in *D. melanogaster* depending on the host’s *dpt* genotype, making it an excellent candidate for examining host genotype impact influences on infection dynamics (Hanson et al., 2019, 2023; Mullinax et al., 2023; Unckless et al., 2015, 2016). Additionally, the genetic diversity within the host population for the *dpt* genotypes, further enriches this model, allowing us to dissect the contributions of specific alleles to infection outcomes (Hanson et al., 2023; Mullinax et al., 2023). The wildtype serine allele (*dpt^S69^*) is associated with higher resistance to *P. rettgeri*, while the segregating arginine allele (*dpt^S69R^*) confers increased susceptibility (Hanson et al., 2023; Mullinax et al., 2023; Unckless et al., 2016). The null allele (Δ*dpt*) results in nearly all flies succumbing to infection within two days, highlighting the crucial role of the *dpt* gene in mediating immune responses (Hanson et al., 2023; Mullinax et al., 2023; Unckless et al., 2016). This variation allows for a robust analysis of how host genetics influence susceptibility and resistance to bacterial infections. Our approach allowed us to vary infection outcomes along three axes: the intrinsic virulence of the pathogen, the dose of the pathogen, and the susceptibility of the host. We then compared pathogenicity metrics across these three axes.

We measured survival against bacterial infections for four different species of *Providencia* injected in variable doses (different ODs) in flies of three different *dpt* genotypes (homozygous serine *dpt* (wildtype, *dpt^S69^*), homozygous arginine *dpt* (*dpt^S69R^*), and *dpt* null (Δ*dpt*)) in a controlled genetic background. The *Providencia* strains we used represent a range from avirulent (*P. burhodogranariea* strain D) to very virulent (*P. alcalifaciens* and *P. sneebia*). The *dpt* genotypes are an allelic series at least in relation to *P. rettgeri* (Mullinax et al., 2023; Unckless et al., 2016). Finally, we considered advantages and limitations of our different pathogenicity measures, offering recommendations on when to use each method of analysis.

## Materials & methods

### Fly stocks and maintenance

We used CRISPR/Cas9 edited flies with modified *diptericin A* gene (Mullinax et al., 2023). Briefly, after editing, flies were isogenized for the X and 3^rd^ chromosomes using a marked strain that allowed us to replace with a specific line’s chromosomes. We did not attempt to do this for the 2^nd^ chromosome. Thus, the second chromosome may contain off-target edits or be an altogether different chromosome if the injected line was segregating for multiple 2^nd^ chromosomes. The latter possibility is unlikely because we Sanger sequenced several immune genes on the 2^nd^ chromosome for our edited lines and all sequences were identical across all the lines. As with any CRISPR/Cas9 editing experiment, we cannot rule out off-target edits, but if they exist, it only slightly changes our interpretation: instead of measuring variation in *dpt* genotype, we are measuring variation in a second chromosome composite genotype. We used three lines of *D. melanogaster,* with the same genetic background, but polymorphic for *diptericin A* gene – homozygous serine *dpt* (wildtype, *dpt^S69^*), homozygous arginine *dpt* (*dpt^S69R^*), and *dpt* null (Δ*dpt,* refers to lines with a 3 base pair deletion). Flies were maintained in a 23°C incubator with a 12h light:12h dark schedule on a cornmeal-molasses-yeast diet (64.3g/L cornmeal, 79.7mL/L molasses, 35.9g/L yeast, 8g/L agar, 15.4mL of food acid mix (50mL Phosphoric Acid + 418mL Propionic Acid + 532mL deionized water) and 1g/L Tegosept.

### Bacterial strains

The following bacteria were used for systemic infections for this study: *Providencia rettgeri* (isolated from wild-caught *D. melanogaster* by B. Lazzaro in State College, PA, USA)*, Providencia burhodograneria* Strain D (isolated from the hemolymph of wild *Drosophila*; P. Juneja and B. Lazzaro), *Providencia sneebia* (isolated from the hemolymph of wild *Drosophila*; P. Juneja and B. Lazzaro), and *Providencia alcalifaciens* (isolated from wild-caught *D. melanogaster* by P. Juneja and S. Short in Ithaca, NY, USA) (Galac & Lazzaro, 2011; Juneja & Lazzaro, 2009; Lazzaro, 2002). Bacteria were grown from glycerol stocks on LB plates at 37°C overnight. Individual colonies of bacteria were picked and grown in 2mL LB broth shaking overnight at 37°C for preparing infection cultures. The optical density (A_600nm_) of the suspension was adjusted according to the desired infection dose.

### Infection assays

We used bacterial suspensions for *P. rettgeri, P. burhodograneria* Strain D, *P. sneebia,* and *P. alcalifaciens* at an Optical Density (A_600nm_) of approximately 0.01, 0.1, 0.5, 1, and 8 for our infections (we use "approximate" concentrations because it is difficult to achieve an exact OD_600_ in bacterial culture preparation, however, the exact concentrations for each treatment are recorded and available in our data file in the column “OD”). We infected a total of 50 flies (10 flies per vial) per genotype per bacteria per optical density. We also infected a total of 50 flies (10 flies per vial) per genotypes with LB media as a control. To induce systemic infection, 5-9 day old flies were pricked in the thorax with a needle dipped in a bacterial suspension. We also used LB media instead of bacterial suspension to poke flies as a control. Flies were incubated post infection at 23°C at a density of 10flies/vial with a 12h light:12h dark schedule and survival was tracked daily for 7 days post infection. The experiments were randomized and conducted across 8 different days.

We used a climbing assay to estimate the fraction of symptomatic (sick) flies following infection, with mortality included as the ultimate symptom. Each day, for seven days post-infection, we assessed the proportion of flies in each vial capable of climbing the vial walls within 5 seconds after being tapped to the bottom. Flies that failed to climb were recorded as symptomatic, and those that had died were also included in the calculation as symptomatic, given that mortality is indicative of severe disease progression. Figure 1A represents a schematic of our experimental design.

**Figure 1:**
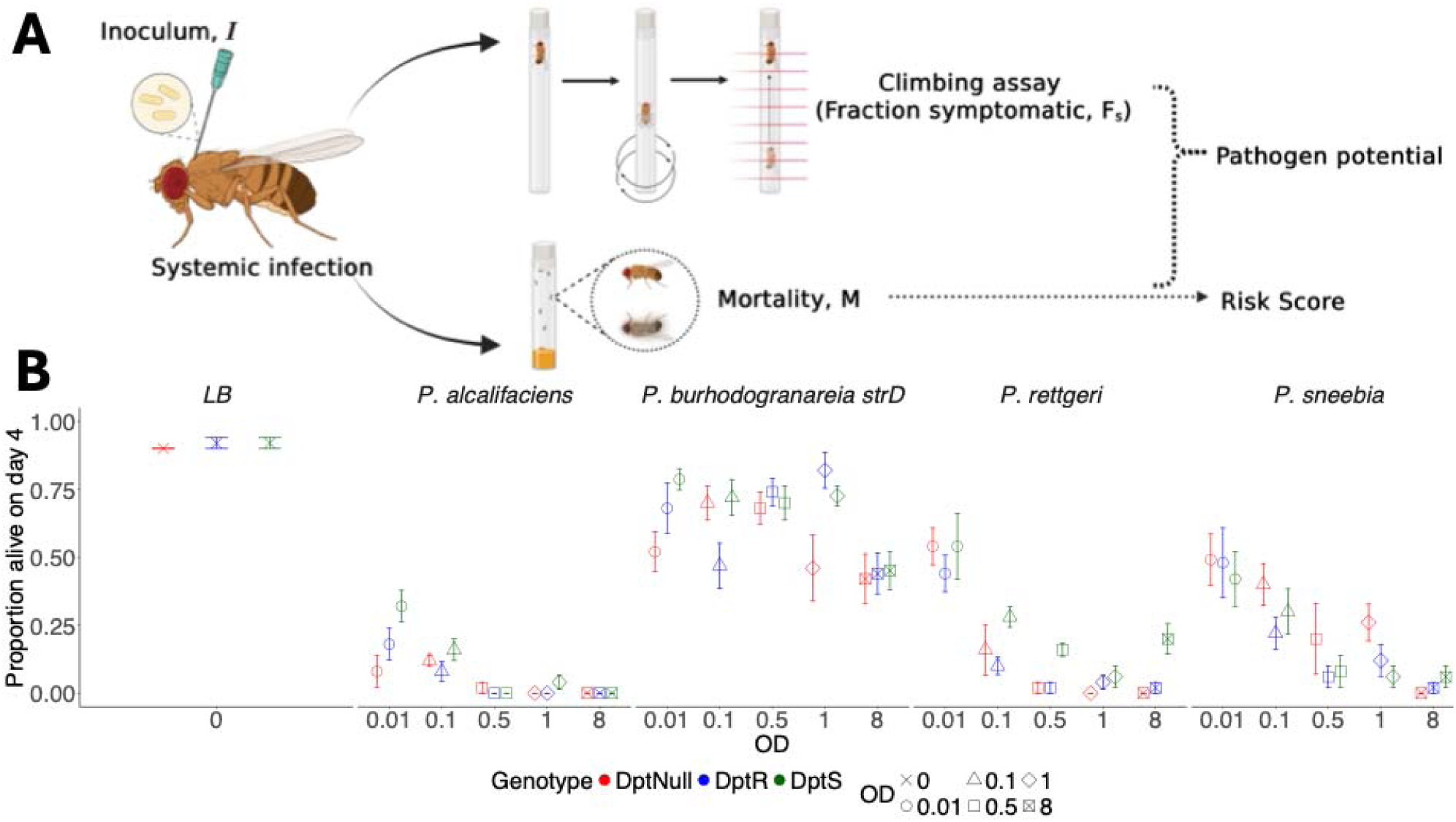
**A.** Experimental design (made using BioRender). 10 flies/vial were infected with variable pathogen strains (*P. burhodograneria* Strain D, *P. rettgeri, P. sneebia, P. alcalifaciens*) with different loads of infective inoculum (OD) and each vial was recorded for upto 7 days post-infection for mortality and fraction symptomatic (using a climbing assay – how many flies can climb up the vial in 5 seconds). Risk scores are further computed using mortality and pathogen potential is calculated using inoculum load, mortality, and fraction symptomatic. **B.** Plot of the proportion of flies alive on day 4 post infection for various host genotypes (homozygous serine *dpt* (wildtype, *dpt^S69^*), homozygous arginine *dpt* (*dpt^S69R^*), and *dpt* null (*Δdpt*)), and variable pathogen strains (*P. burhodograneria* Strain D, *P. rettgeri, P. sneebia, P. alcalifaciens*) with different loads of infective inoculum (OD). The error bars represent the 95% confidence interval around the mean of the proportion of flies alive on day 4 post infection.

### Bacterial load

To measure actual dosing in newly infected flies, single wildtype, *dpt^S69^* flies were pricked using *P. rettgeri* and *P. sneebia* cultures at an optical density (A_600nm_) of approximately 0.01, 0.1, 0.5, 1, and 8, (exact concentrations for each treatment are recorded and available in our data file in the column “OD”) followed by surface sterilization by washing in 70% ethanol and molecular grade water, and then, homogenized in 500μL LB and plated on LB plates immediately after the infection using a WASP touch spiral plater from Don Whitely Scientific. The LB agar plates were incubated overnight at 37°C. Gut commensal bacteria grow more slowly than *Providencia* under these conditions, so by limiting incubation to overnight we exclude any commensal bacteria from our assay. The number of colony forming units (CFU) on each plate was recorded using a Neutec Flash and Go colony counter. We used 5 replicates (using single flies) per bacteria per A_600nm_ for quantifying the systemic bacterial load in infected flies. We chose to measure bacterial load for only two species in this study (*P. rettgeri* and *P. sneebia*) because, based on previous studies (Beal et al., 2020), it is established that bacterial load does not vary significantly across species within the same genus at the same optical densities (ODs). As a result, we did not perform bacterial load assays for the other two pathogens, *P. burhodograneria* Strain D and *P. alcalifaciens*, as we did not expect different outcomes at equivalent ODs.

### Data and statistical analysis

We fitted our data to a mixed effects cox regression model to analyze the effect of the type and dose of pathogen and the host genotype. We performed a principal component analysis and a correlation matrix to look at the variation between the different measures of virulence and pathogenicity. Data analysis performed done in R version 4.3.1 (R Core Team, 2013). Data visualization was performed using R packages – ggplot2 (Wickham, 2011), survminer (Kassambara et al., 2017), wesanderson (Ram & Wickham, 2018), ggpubr (Kassambara & Kassambara, 2020), corrplot (Wei et al., 2017), RColorBrewer (Neuwirth & Neuwirth, 2014) and ggrepel (Slowikowski et al., 2018). Data was fitted to a cox-proportional hazards model to analyze the effect of host genotype, pathogen type and dose on survival [Table S2]. Statistical analysis was performed using R packages – lme4 (Bates, 2010) and coxme (Therneau & Therneau, 2015). We have included date as a random factor in all statistical analyses to account for variability across experimental days.

## Results

### Host genotype, dose and pathogen strain interact to influence survival against infection

We measured survival against bacterial infections for four different species of *Providencia* injected in variable doses (different ODs) in flies of three different *dpt* genotypes (homozygous serine *dpt* (wildtype, *dpt^S69^*), homozygous arginine *dpt* (*dpt^S69R^*), and *dpt* null (Δ*dpt*)) in a controlled genetic background. These *Providencia* strains represent a range from avirulent (*P. burhodogranereia*) to highly virulent (*P. alcalifaciens* and *P. sneebia*) bacteria (Galac & Lazzaro, 2011). The host *dpt* genotypes are an allelic series at least in relation to *P. rettgeri* where the wildtype serine allele is most resistant, the segregating arginine allele is more susceptible, and the null allele results in almost all flies dying within two days of infection (Mullinax et al., 2023). Our results from our infection experiments align with previously published findings - (i) flies homozygous for the serine allele survive systemic infection with *P. rettgeri* much better than those homozygous for the arginine allele (Unckless et al., 2016), (ii) *P. alcalifaciens* and *P. sneebia* are highly pathogenic to *D. melanogaster*, where small doses of infection with these bacteria can result in lethal infections (Galac & Lazzaro, 2011), (iii) most flies survive infections against *P. burhodograneria* Strain D at moderate doses (Galac & Lazzaro, 2011) [Figure 1B, Figure 2A].

**Figure 2:**
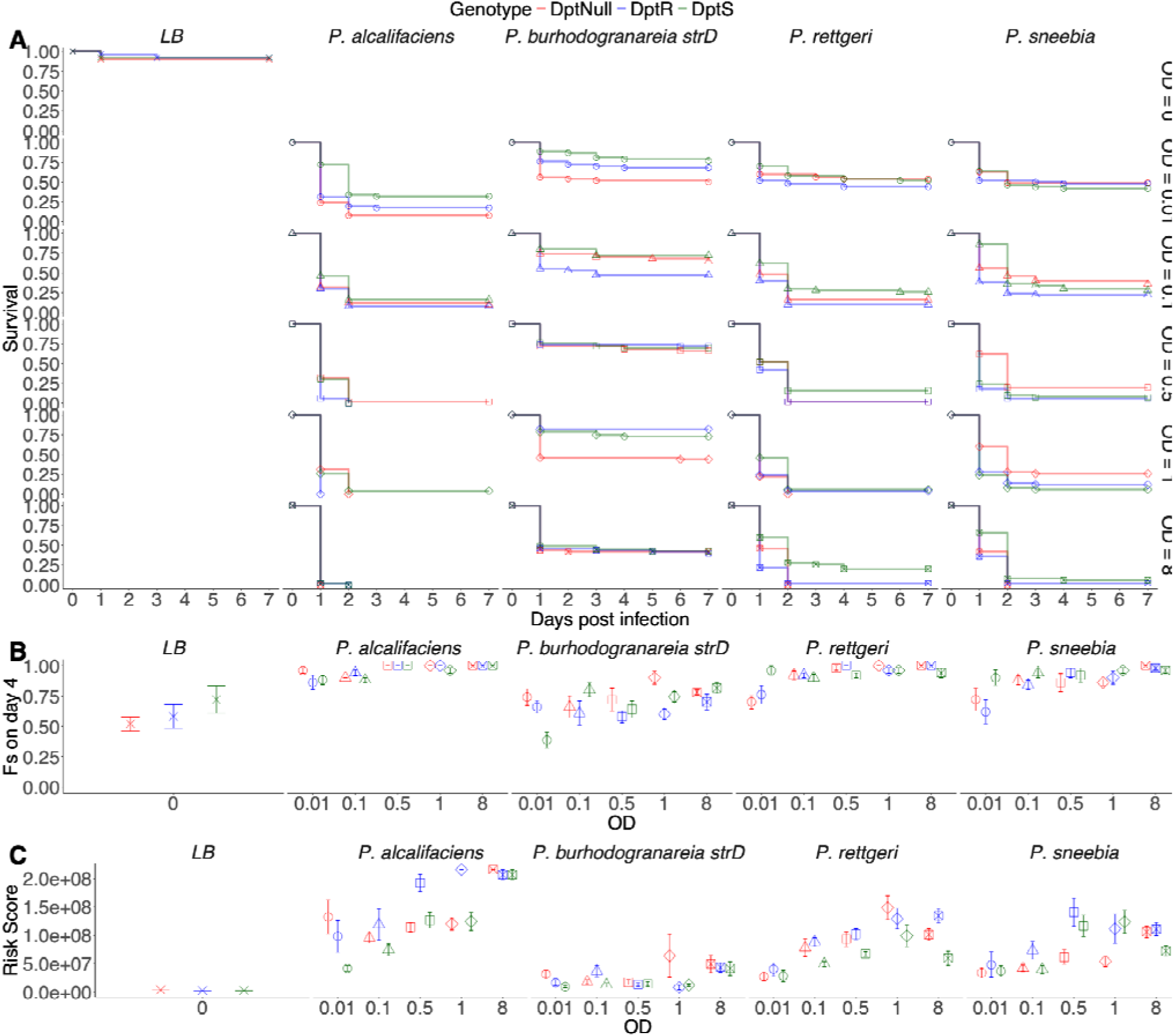
Host-based measures of pathogenicity: **A.** Survival curves show proportion alive 0-7 days post infection in various host genotypes (homozygous serine *dpt* (wildtype, *dpt^S69^*), homozygous arginine *dpt* (*dpt^S69R^*), and *dpt* null (*Δdpt*)), and variable pathogen strains (*P. burhodograneria* Strain D, *P. rettgeri, P. sneebia, P. alcalifaciens*) with different loads of infective inoculum (OD) ranging from OD_600_ = 0-8. **B.** Fraction symptomatic four days post-infection in the same assays as A calculated following climbing assays. **C.** Risk scores for the same assays as A calculated based on the Cox models.

We surveyed survival against infection using Kaplan-Meier survival curves and proportion alive on day 4 post-infection to assess the effect of host genotype, dose, and pathogen strain [Figure 1B, Figure 2A]. We rationalized that 4 days post infection was a reasonable snapshot because most flies destined to die had died by that point so the interaction between host immunity and pathogen was mostly over. We calculated the fraction of flies that died between day 4 and day 7 post-infection to justify this timepoint and only 2% of flies die between day 4 and day 7 post-infection. Very small doses of *P. alcalifaciens* were sufficient to cause high mortality in flies regardless of their genotype. We saw a range of rates of mortality for survival against *P. sneebia* across genotypes and across different doses of infection. While for survival against *P. burhodograneria* Strain D, even high doses of injection did not prove to be lethal for most of the flies. We also quantified the bacterial load immediately after infection for the range of doses (optical densities) used in our assays. Dose was not linear with the optical density, instead the optical density was linearly related to log transformation of the dose (Figure S1, Table S1). We looked at the effect of host genotype, pathogen type and dose on survival using a cox-proportional hazards model [Table S2]. First, both the dose (OD) and pathogen species significantly affect survival, with *P. alcalifaciens* showing the highest pathogenicity (Table S2: coef = 1.273, p < 0.001). Second, host genotype markedly influences survival outcomes: Δ*dpt* flies showed a higher mortality against all pathogens except *P. sneebia*, while the genotypic difference between *dpt^S69^*and *dpt^S69R^* did not significantly influence survival against the different pathogen strains except for *P. rettgeri* where flies with the serine allele (*dpt^S69^*) have significantly better survival rates compared to those with the arginine allele (*dpt^S69R^*) or null allele (Δ*dpt*) (Table S2: GenotypeDptS: coef = -1.116, p = 0.001). Third, interactions between OD, treatment, and genotype underscore the complexity of these relationships, highlighting specific combinations that significantly alter survival rates (Table S2: OD1 * Treatment*P. rettgeri* * GenotypeDptS (coef = -1.566, p = 0.005)).

### Limitations of measures of pathogenicity

#### Host-based measures

We examined Kaplan-Meier survival curves, risk scores, and fraction symptomatic to assess host-based measures of pathogenicity [Figure 2]. Both risk scores and fraction symptomatic follow an approximately inverse relationship to host survival upon infection [Figure 2]. Fraction symptomatic is least sensitive to host genotype out of the three measures of host susceptibility [Figure 2]. Risk scores show variation with both the dose and virulence of pathogens but fail to resolve the interaction between the dose of a pathogen and the host genotype [Figure 2]. These host-based measures of pathogenicity are easy to interpret, but they tell us little about pathogen virulence of individual microbes. It becomes difficult to determine whether the differences in survival seen in the host is due to a highly virulent pathogen or infection with a high dose of a pathogen. Most pathogens become virulent enough to cause visible symptoms/mortality when administered to a single host at high doses.

### Pathogen-based measures

We looked at the pathogen potential and related metrics to assess pathogen-based measures of pathogenicity [Figure 3]. We quantified the pathogen potential on day 4 post-infection, and on the day of median survival for each vial. Pathogen potential was proposed as a tool for adding quantitative rigor to comparisons in microbial pathogenesis (Casadevall, 2017, 2022; Casadevall et al., 2011; Casadevall & Pirofski, 1999, 2001, 2003; Smith & Casadevall, 2022). Pathogen potential is helpful in analyzing the dose-dependent effect of pathogen on host mortality, but there are limitations to the measure. Pathogen potential failed to capture the variation between host genotypes [Figure 4A]. The limitation in assessing pathogen potential arises when evaluating high doses or highly virulent microbes at moderate doses, wherein the likelihood of a specific pathogenic microbe being deemed the primary cause of mortality approaches zero, even though all individuals die. This counterintuitive phenomenon stems from the low probability that any specific pathogen cell within the high dose is the sole contributor to mortality. While this concept aligns with the understanding that the pathogen responsible for mortality may not be easily pinpointed, it presents a challenge in practical utility in our context, as the assessment becomes less informative with more virulent pathogens.

**Figure 3:**
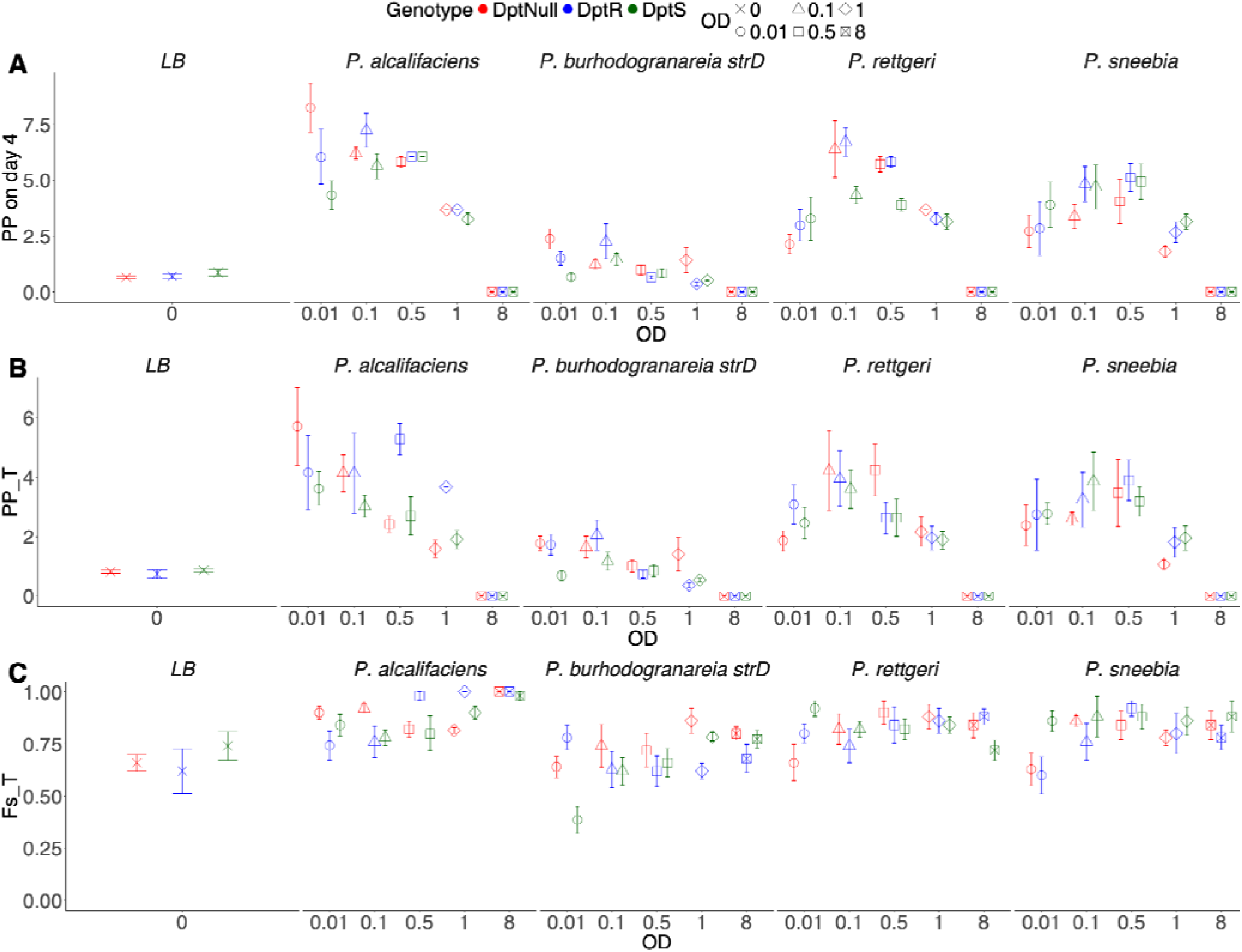
Pathogen-based measures of pathogenicity: **A.** Pathogen potential four days post-infection calculated based on the inoculum load, host mortality and fraction symptomatic for various host genotypes (homozygous serine *dpt* (wildtype, *dpt^S69^*), homozygous arginine *dpt* (*dpt^S69R^*), and *dpt* null (*Δdpt*)), and variable pathogen strains (*P. burhodograneria* Strain D, *P. rettgeri, P. sneebia, P. alcalifaciens*) with different loads of infective inoculum (OD) ranging from OD_600_ = 0-8. **B.** Pathogen potential on median survival day (T) calculated based on the inoculum load, host mortality, fraction symptomatic and day post-infection for the same assays as A. **C.** Fraction symptomatic on median survival day (T) for the same assays as A calculated following climbing assays.

**Figure 4:**
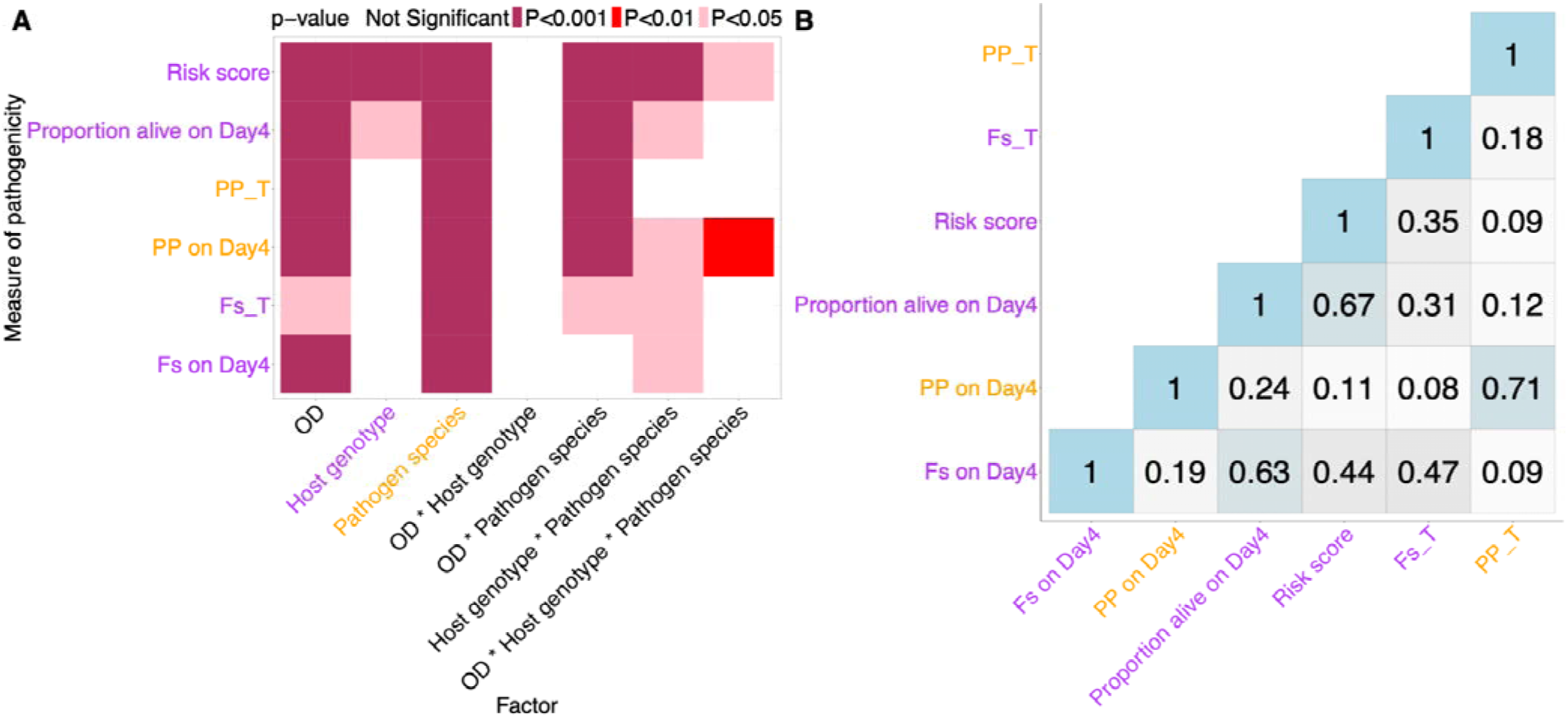
A. P-values from ANOVA to indicate the significance of each factor corresponding to each measure of pathogenicity (see Table S3 for more information), B. Heatmap showing R^2^ (R=correlation coefficient) values to assess the differences between different measures of host susceptibility (survival, risk scores, and fraction symptomatic) and pathogen virulence (pathogen potential) (see Table S4 for more information). [Host-based measures are labelled purple while pathogen-based measures are labelled orange]

In this context, it is important to consider the transparency of the CRISPR methodology employed in these experiments. Specifically, the application of CRISPR-Cas9 in generating the *dpt* mutant strains (Δ*dpt*, *dpt^S69^*, and *dpt^S69R^*) could introduce concerns about off-target effects, particularly since chromosome 2, which houses many immune-related genes, was not backcrossed. We acknowledge the possibility of off-target effects, particularly considering the sensitivity of the *dpt^S69R^* mutant to Gram-positive bacteria, a pathogen type that *dpt* does not typically impact (Carboni et al., 2022; Hanson et al., 2019). To address these concerns, we conducted Sanger sequencing to evaluate potential off-target effects. While we cannot entirely exclude such effects, our data suggest they are minimal. We opted against extensive backcrossing due to potential confounding genetic variability and focused instead on robust sequencing to reduce possible confounders. Importantly, even if off-target effects are present, each of our genotypes represents a distinct chromosome 2 background. Thus, we are not attributing effects to a single mutation but rather assessing the cumulative impact of the entire second chromosome.

### Comparative analysis

We took a comparative approach to determine how different measures of host susceptibility (survival, risk scores, and fraction symptomatic) and pathogen virulence (pathogen potential) were correlated with each other to assess how different they are in their analytical information. From the R^2^ matrix, it was evident that all the different host-based measures of virulence and susceptibility were highly correlated to each other, but not to the pathogen-based measures (pathogen potential) [Figure 4B]. To further examine the relationship among metrics, we performed a principal component analysis (PCA). This revealed three main groupings: risk score (alone), fraction alive and fraction symptomatic metrics, and pathogen-potential related metrics [Figure S2]. Pathogen potential metrics clustered away from the host-based measures as they incorporate quantitative information about the infecting pathogen as well as host response. Interestingly, risk scores also clustered away from all the different pathogenicity measures as risk scores were able to capture the information for the variation between host genotype, pathogen type and dose of the pathogen [Figure 4A, Figure S2].

## Discussion

Our study investigated the complex dynamics of host-pathogen interactions, focusing on the quantification of host susceptibility and pathogenicity. Traditionally, host mortality rates, analyzed through the Cox proportional hazards models, have been widely used to assess virulence. An alternative, pathogenic potential concept, introduced by Casadevall, 2017 offers a more nuanced perspective by considering the composite nature of host damage. In this study, we explore the advantages and limitations of traditional survival analysis and risk scores compared to the quantitative measures like pathogen potential. Our results suggest that, at least for our study system, the best measure depends on overall host mortality in relation to dose.

Our infection experiments revealed intricate interactions between host genotype, pathogen strain, and infection dose [Table S2]. First, in most cases, mortality increased with increasing dose and this was reflected in most of our measures of infection outcomes. However, because pathogen potential (and related metric) explicitly control for the effect of dose, those measures are less sensitive to dose variation. Second, host genotype influences the outcomes of infection for most, but not all infections and infection outcome measures. The observed higher mortality in Δ*dpt* flies against all pathogens, except *P. sneebia*, underscores the role of the diptericin A gene in systemic infection resilience (Hanson et al., 2023; Mullinax et al., 2023; Unckless et al., 2016). Finally, pathogen species consistently displayed variation in pathogenicity (*sensu latu*) across infection outcome measures. *P. alcalifaciens* demonstrated high pathogenicity even at low doses, emphasizing the importance of considering both pathogen virulence and infection dose in assessments.

Traditional survival analysis and risk scores, rooted in Cox models, proved informative for overall trends and risk assessment. However, they fall short in providing a comprehensive understanding of pathogen virulence, particularly for avirulent or less virulent pathogens. The limitations of these host-based measures highlight the necessity for additional quantitative insights. The pathogen-based measure, pathogen potential, offers a quantitative lens but encounters limitations at high pathogen doses, where pinpointing the primary cause of mortality becomes less informative. It becomes evident that each measure contributes unique insights, emphasizing the importance of a nuanced and multifaceted approach in analyzing host-pathogen interactions. A combination of pathogen potential and risk scores seemed to be particularly informative in separating the variation between host susceptibility and pathogen virulence.

Additionally, concepts such as Systemic Pathogen Burden Load (SPBL) and Bacterial Load Undergoing Dynamics (BLUD) could further enrich the biological context of the manifestation of symptoms during infection (Duneau et al., 2017). These measures, which focus on pathogen load and its dynamics over time, offer insights into infection progression and immune response that complement the survival data and risk scores we used. Although our current study does not include these measures, this could provide a more detailed understanding of immune responses in the early phases of infection, which aligns with recent work on time-specific infection dynamics.

We propose several recommendations for the optimal use of each measure, considering their strengths and limitations –

i. <u>Kaplan-Meier Curves (to visualize survival functions):</u> Kaplan-Meier curves are a robust measure and should be used to assess the overall survival trends over time. It is particularly useful for understanding the impact of host genotype, dose, and pathogen strain on the course of infection. **Limitations:** While survival analysis provides valuable information about the overall outcome, it may not distinguish between the effects of pathogen virulence and the dose administered.
ii. <u>Risk Scores from Cox Proportional Hazards Models:</u> Risk scores derived from Cox proportional hazards models are informative for assessing variation in both dose and virulence of pathogens. They are especially useful when trying to understand how different combinations of host genotype, dose, and pathogen strain contribute to the risk of mortality. **Limitations:** Risk scores may not provide information about the virulence of avirulent or less virulent pathogens, and their utility diminishes for high doses of less virulent pathogens.
iii. <u>Fraction Symptomatic:</u> Fraction symptomatic is useful for insights into the proportion of individuals exhibiting symptoms, giving a qualitative assessment of the infection. **Limitations:** Fraction symptomatic is less sensitive to pathogen virulence and may not provide detailed information about the severity of the infection.
iv. <u>Pathogen Potential:</u> Pathogen potential is valuable for understanding the dose-dependent effects of pathogens on host mortality. **Limitations:** Pathogen potential may be less informative when mortality is high since it loses resolution, where the likelihood of pinpointing a specific pathogen as the primary cause of mortality decreases.

Survival analysis is robust for overall trends, risk scores offer detailed information on variation, fraction symptomatic provides qualitative insights, and pathogen potential contributes quantitative rigor. A comparative analysis aids in choosing the most relevant measure based on experimental goals.

## Acknowledgements

We thank Brian Lazzaro for providing the *Providencia* strains, and Unckless lab members for helpful comments and advice. We also thank two anonymous reviewers who provided feedback that ultimately greatly improved the manuscript. The work was supported by NSF 2330095 and NIH R01-AI139154 to RLU and funds from KU Entomology Endowment to AG.

## Data accessibility

Data and code is archived on Zenodo/GitHub (https://doi.org/10.5281/zenodo.15116399 / https://github.com/anjaligupta1210/Pathogenicity-across-variation-in-microbial-virulence-and-host-resistance.git).

