## Supplementary Information for "On pathogenicity with varying microbial virulence and host resistance"

**Figure S1:** Bacterial load (CFU/fly) vs optical density (A600nm) for *P. rettgeri* and *P. sneebia*. Single wildtype *dpt^S69^* flies were infected with bacteria at OD_600_ 0–8, surface sterilized, homogenized in 500 µL LB, and plated immediately using a spiral plater. Plates were incubated overnight at 37°C followed by counting colony forming units (CFUs). Five single-fly replicates were used per OD per pathogen species. Statistical analysis can be seen in Table S1.

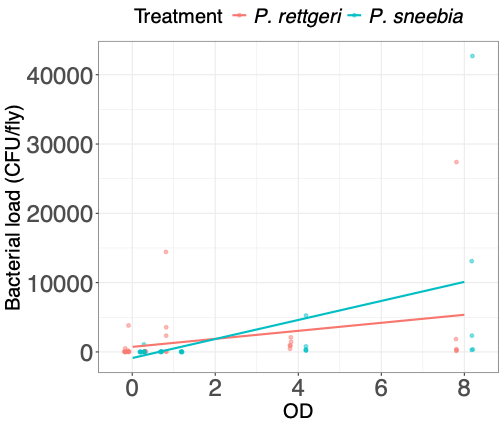

**Figure S2:** Principal component analysis (PCA) over different measures of host susceptibility (survival, risk scores, and fraction symptomatic) and pathogen virulence (pathogen potential). [Prop_DayX_Alive: Proportion survival on Day X post-infection, PP_DayX: Pathogen potential on Day X post-infection, PP_T: Pathogen potential on median survival day, F_s__T/T: Fraction symptomatic on median survival day/ Median survival day, Fs_DayX: Fraction symptomatic on Day X post-infection, Fs_T: Fraction symptomatic on median survival day].

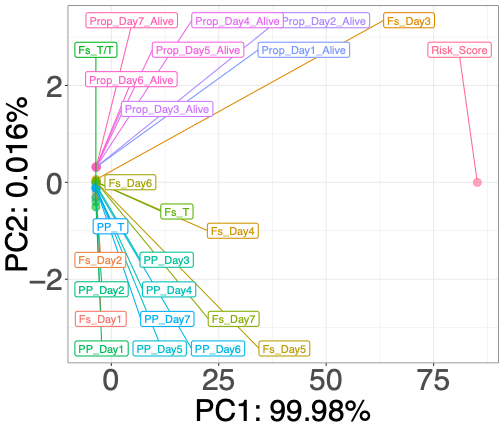

**Table S1:** Statistical analysis for analyzing the effect of OD and treatment (pathogen species) on log-transformed bacterial load. Data was fitted to a linear mixed-effects model.

[Model: lm((`Bacterial_Load_CFU/fly`+1) ~ OD + Treatment)]

|  | Estimate | Std. Error | t value | Pr(>\|t\|) |
| --- | --- | --- | --- | --- |
| (Intercept) | -173.1791 | 1319.68112 | -0.131228 | 0.89605704 |
| OD | 976.346408 | 281.732659 | 3.46550667 | 0.00101252 |
| Treatment_P. sneebia | 191.833333 | 1632.88456 | 0.11748126 | 0.90689169 |

**Table S2:** Statistical analysis for analyzing the effect of OD, treatment (pathogen species), and host genotype on survival. Data was fitted to a mixed-effects cox-proportional hazards model.

[Model: coxph(Survival ~ OD * Treatment * Genotype + frailty(Date)]

|  | coef | se(coef) | se2 | Chisq | DF | p | significance level |
| --- | --- | --- | --- | --- | --- | --- | --- |
| OD0.1 | 0.30174574 | 0.26403205 | 0.26403205 | 1.30607764 | 1 | 0.25310605 | Not significant |
| OD0.5 | 0.62891245 | 0.25203071 | 0.25202873 | 6.22692227 | 1 | 0.01258222 | * |
| OD1 | 0.51912021 | 0.25596387 | 0.25596191 | 4.11318777 | 1 | 0.04255009 | * |
| OD8 | 1.20718935 | 0.24229568 | 0.24229568 | 24.8232953 | 1 | 6.28E-07 | *** |
| GnDptR | 0.08505831 | 0.27735556 | 0.27735556 | 0.0940502 | 1 | 0.75908996 | Not significant |
| GnDptS | 0.1402146 | 0.27008436 | 0.27008436 | 0.26951788 | 1 | 0.60365536 | Not significant |
| TrtP. alcalifaciens | 1.27732984 | 0.24571707 | 0.24571504 | 27.0231176 | 1 | 2.01E-07 | *** |
| TrtP. burhodogranareia strD | 0.00391368 | 0.28011647 | 0.28011468 | 0.00019521 | 1 | 0.98885262 | Not significant |
| TrtP. rettgeri | -0.1366926 | 0.28626178 | 0.28626004 | 0.22801498 | 1 | 0.63299961 | Not significant |
| frailty(Date) | NA | NA | NA | 0.0004676 | 3.40E-05 | 0.86358273 | Not significant |
| OD0.1:GnDptR | 0.41437501 | 0.36585899 | 0.36585899 | 1.28280205 | 1 | 0.25737869 | Not significant |
| OD0.5:GnDptR | 0.6967196 | 0.35114324 | 0.35114183 | 3.93683698 | 1 | 0.04723952 | * |
| OD1:GnDptR | 0.51866575 | 0.35597611 | 0.35597583 | 2.12291904 | 1 | 0.14510975 | Not significant |
| OD8:GnDptR | -0.0501395 | 0.34253726 | 0.34253726 | 0.02142613 | 1 | 0.88362405 | Not significant |
| OD0.1:GnDptS | -0.1813748 | 0.36330025 | 0.36330025 | 0.24924284 | 1 | 0.61760873 | Not significant |
| OD0.5:GnDptS | 0.49145382 | 0.34602362 | 0.34602323 | 2.01722309 | 1 | 0.15552333 | Not significant |
| OD1:GnDptS | 0.6442761 | 0.34831348 | 0.34831312 | 3.42139697 | 1 | 0.06435653 | Not significant |
| OD8:GnDptS | -0.5103881 | 0.33801418 | 0.33801418 | 2.27998143 | 1 | 0.13105344 | Not significant |
| OD0.1:TrtP. alcalifaciens | -0.4679878 | 0.3371746 | 0.3371746 | 1.92645756 | 1 | 0.16514653 | Not significant |
| OD0.5:TrtP. alcalifaciens | -0.611933 | 0.32500924 | 0.3250077 | 3.54500117 | 1 | 0.05972512 | Not significant |
| OD1:TrtP. alcalifaciens | -0.450733 | 0.32873275 | 0.32873123 | 1.8799783 | 1 | 0.17033661 | Not significant |
| OD8:TrtP. alcalifaciens | -0.5733042 | 0.31677745 | 0.31677587 | 3.27538062 | 1 | 0.07032661 | Not significant |
| OD0.1:TrtP. burhodogranareia strD | -0.8439314 | 0.40654376 | 0.40654376 | 4.30923015 | 1 | 0.03790611 | * |
| OD0.5:TrtP. burhodogranareia strD | -1.2098992 | 0.40296058 | 0.40295933 | 9.01515699 | 1 | 0.0026775 | ** |
| OD1:TrtP. burhodogranareia strD | -0.283935 | 0.37580038 | 0.37579905 | 0.57085199 | 1 | 0.44992056 | Not significant |
| OD8:TrtP. burhodogranareia strD | -0.8951119 | 0.36488414 | 0.36488414 | 6.01789621 | 1 | 0.01416152 | * |
| OD0.1:TrtP. rettgeri | 0.66829828 | 0.37027156 | 0.37027021 | 3.25761673 | 1 | 0.07109238 | Not significant |
| OD0.5:TrtP. rettgeri | 0.56469765 | 0.35696996 | 0.35696717 | 2.50246845 | 1 | 0.11366801 | Not significant |
| OD1:TrtP. rettgeri | 1.10008229 | 0.3594512 | 0.35944842 | 9.36635209 | 1 | 0.00221005 | ** |
| OD8:TrtP. rettgeri | 0.08968093 | 0.34920605 | 0.34920605 | 0.06595332 | 1 | 0.79732253 | Not significant |
| GnDptR:TrtP. alcalifaciens | -0.3983181 | 0.34999305 | 0.34999305 | 1.29521337 | 1 | 0.25508941 | Not significant |
| GnDptS:TrtP. alcalifaciens | -1.1104188 | 0.35244436 | 0.35244294 | 9.92641703 | 1 | 0.00162923 | ** |
| GnDptR:TrtP. burhodogranareia strD | -0.7495597 | 0.4235912 | 0.4235912 | 3.13125638 | 1 | 0.07680453 | Not significant |
| GnDptS:TrtP. burhodogranareia strD | -1.2571298 | 0.4430512 | 0.4430512 | 8.05104754 | 1 | 0.00454774 | ** |
| GnDptR:TrtP. rettgeri | 0.23298565 | 0.39512658 | 0.39512658 | 0.34768492 | 1 | 0.55542658 | Not significant |
| GnDptS:TrtP. rettgeri | -0.1566122 | 0.39760536 | 0.39760536 | 0.15514817 | 1 | 0.69366334 | Not significant |
| OD0.1:GnDptR:TrtP. alcalifaciens | -0.0156989 | 0.47257479 | 0.47257479 | 0.00110356 | 1 | 0.97349924 | Not significant |
| OD0.5:GnDptR:TrtP. alcalifaciens | 0.09718381 | 0.45736296 | 0.45736186 | 0.04515084 | 1 | 0.8317269 | Not significant |
| OD1:GnDptR:TrtP. alcalifaciens | 0.36009204 | 0.46174233 | 0.46174211 | 0.60817372 | 1 | 0.43547627 | Not significant |
| OD8:GnDptR:TrtP. alcalifaciens | 0.31588541 | 0.45044083 | 0.45044083 | 0.49179447 | 1 | 0.48312792 | Not significant |
| OD0.1:GnDptS:TrtP. alcalifaciens | 0.89375434 | 0.47876579 | 0.47876579 | 3.48489834 | 1 | 0.06193117 | Not significant |
| OD0.5:GnDptS:TrtP. alcalifaciens | 0.54758607 | 0.45965657 | 0.45965627 | 1.41918115 | 1 | 0.23353823 | Not significant |
| OD1:GnDptS:TrtP. alcalifaciens | 0.31546358 | 0.46267744 | 0.46267716 | 0.46488101 | 1 | 0.49535162 | Not significant |
| OD8:GnDptS:TrtP. alcalifaciens | 1.43307852 | 0.45342293 | 0.45342183 | 9.98925305 | 1 | 0.00157456 | ** |
| OD0.1:GnDptR:TrtP. burhodogranareia strD | 0.868055 | 0.57351035 | 0.57351035 | 2.29093073 | 1 | 0.13013187 | Not significant |
| OD0.5:GnDptR:TrtP. burhodogranareia strD | -0.2491324 | 0.59667312 | 0.59667228 | 0.17433613 | 1 | 0.67628656 | Not significant |
| OD1:GnDptR:TrtP. burhodogranareia strD | -1.4085857 | 0.61325413 | 0.61325397 | 5.27576714 | 1 | 0.02162426 | * |
| OD8:GnDptR:TrtP. burhodogranareia strD | 0.70993474 | 0.5363294 | 0.5363294 | 1.75215957 | 1 | 0.18560548 | Not significant |
| OD0.1:GnDptS:TrtP. burhodogranareia strD | 0.98972251 | 0.6183106 | 0.6183106 | 2.56220291 | 1 | 0.10944598 | Not significant |
| OD0.5:GnDptS:TrtP. burhodogranareia strD | 0.46157714 | 0.60706591 | 0.60706568 | 0.57811854 | 1 | 0.44705074 | Not significant |
| OD1:GnDptS:TrtP. burhodogranareia strD | -0.6206348 | 0.5931367 | 0.59313649 | 1.09487039 | 1 | 0.2953946 | Not significant |
| OD8:GnDptS:TrtP. burhodogranareia strD | 1.53408453 | 0.55481096 | 0.55481096 | 7.64554629 | 1 | 0.00569127 | ** |
| OD0.1:GnDptR:TrtP. rettgeri | -0.530469 | 0.5089883 | 0.5089883 | 1.08618667 | 1 | 0.29731767 | Not significant |
| OD0.5:GnDptR:TrtP. rettgeri | -0.9030956 | 0.49327302 | 0.49327201 | 3.35191261 | 1 | 0.06712698 | Not significant |
| OD1:GnDptR:TrtP. rettgeri | -0.9550582 | 0.49681353 | 0.49681334 | 3.6954967 | 1 | 0.05455954 | Not significant |
| OD8:GnDptR:TrtP. rettgeri | 0.01006153 | 0.48678236 | 0.48678236 | 0.00042723 | 1 | 0.98350932 | Not significant |
| OD0.1:GnDptS:TrtP. rettgeri | -0.1444547 | 0.51770238 | 0.51770238 | 0.07785798 | 1 | 0.78022147 | Not significant |
| OD0.5:GnDptS:TrtP. rettgeri | -0.7372941 | 0.49915894 | 0.49915867 | 2.18174434 | 1 | 0.1396561 | Not significant |
| OD1:GnDptS:TrtP. rettgeri | -1.0638774 | 0.4979131 | 0.49791285 | 4.56537066 | 1 | 0.03262461 | * |
| OD8:GnDptS:TrtP. rettgeri | -0.0322691 | 0.49437598 | 0.49437598 | 0.00426048 | 1 | 0.94795716 | Not significant |

**Table S3:** ANOVA for analyzing the effect of OD, treatment (pathogen species), and host genotype on different measures of pathogenicity, corresponding to Figure 4A.

[Model: aov(“Measure of pathogenicity” ~ OD * Treatment * Genotype + Error(Vial_ID, Date)]

A. **Risk score**

|  | Df | Sum Sq | Mean Sq | F value | Pr(>F) |
| --- | --- | --- | --- | --- | --- |
| OD | 1 | 1.45E+17 | 1.45E+17 | 139.439144 | 1.97E-26 |
| Genotype | 2 | 2.25E+16 | 1.12E+16 | 10.784722 | 3.06E-05 |
| Treatment | 1 | 9.19E+16 | 9.19E+16 | 88.2284336 | 2.03E-18 |
| OD:Genotype | 2 | 8.60E+14 | 4.30E+14 | 0.41292323 | 0.66211111 |
| OD:Treatment | 3 | 5.72E+16 | 1.91E+16 | 18.3157708 | 6.88E-11 |
| Genotype:Treatment | 8 | 3.99E+16 | 4.99E+15 | 4.78744988 | 1.55E-05 |
| OD:Genotype:Treatment | 6 | 1.43E+16 | 2.38E+15 | 2.2865779 | 0.03586364 |
| Residuals | 283 | 2.95E+17 | 1.04E+15 | NA | NA |

B. **Proportion alive on day 4**

|  | Df | Sum Sq | Mean Sq | F value | Pr(>F) |
| --- | --- | --- | --- | --- | --- |
| OD | 1 | 1.85590565 | 1.85590565 | 92.5478399 | 3.87E-19 |
| Genotype | 2 | 0.15274083 | 0.07637041 | 3.80833846 | 0.02333057 |
| Treatment | 1 | 5.66537229 | 5.66537229 | 282.513267 | 1.91E-44 |
| OD:Genotype | 2 | 0.004357 | 0.0021785 | 0.10863443 | 0.89709569 |
| OD:Treatment | 3 | 0.41292145 | 0.13764048 | 6.86367303 | 0.00017709 |
| Genotype:Treatment | 8 | 0.3470236 | 0.04337795 | 2.16311406 | 0.03039029 |
| OD:Genotype:Treatment | 6 | 0.11755471 | 0.01959245 | 0.9770104 | 0.44100379 |
| Residuals | 283 | 5.67513297 | 0.02005347 | NA | NA |

C. **Pathogen potential on median survival day (PP_T)**

|  | Df | Sum Sq | Mean Sq | F value | Pr(>F) |
| --- | --- | --- | --- | --- | --- |
| OD | 1 | 141.04298 | 141.04298 | 82.2680445 | 2.05E-17 |
| Genotype | 2 | 0.59682808 | 0.29841404 | 0.17405999 | 0.84033638 |
| Treatment | 1 | 41.6897681 | 41.6897681 | 24.3169543 | 1.39E-06 |
| OD:Genotype | 2 | 0.1696832 | 0.0848416 | 0.04948671 | 0.95172604 |
| OD:Treatment | 3 | 43.94572 | 14.6485733 | 8.54427128 | 1.89E-05 |
| Genotype:Treatment | 8 | 16.7925766 | 2.09907208 | 1.22435413 | 0.2844434 |
| OD:Genotype:Treatment | 6 | 11.1680146 | 1.86133576 | 1.08568646 | 0.3710431 |
| Residuals | 283 | 485.184297 | 1.71443215 | NA | NA |

D. **Pathogen potential on day 4**

|  | Df | Sum Sq | Mean Sq | F value | Pr(>F) |
| --- | --- | --- | --- | --- | --- |
| OD | 1 | 314.384138 | 314.384138 | 192.030114 | 1.10E-33 |
| Genotype | 2 | 1.2153476 | 0.6076738 | 0.37117544 | 0.69025826 |
| Treatment | 1 | 102.757145 | 102.757145 | 62.7654639 | 5.37E-14 |
| OD:Genotype | 2 | 0.38239344 | 0.19119672 | 0.11678556 | 0.88981883 |
| OD:Treatment | 3 | 91.8718292 | 30.6239431 | 18.7055216 | 4.26E-11 |
| Genotype:Treatment | 8 | 33.4407918 | 4.18009898 | 2.5532614 | 0.01060474 |
| OD:Genotype:Treatment | 6 | 37.9589005 | 6.32648342 | 3.86430225 | 0.00100704 |
| Residuals | 283 | 463.316452 | 1.63716061 | NA | NA |

E. **Fraction symptomatic on median survival day (F_S__T)**

|  | Df | Sum Sq | Mean Sq | F value | Pr(>F) |
| --- | --- | --- | --- | --- | --- |
| OD | 1 | 0.12610385 | 0.12610385 | 6.35103611 | 0.01228162 |
| Genotype | 2 | 0.06123571 | 0.03061786 | 1.54202354 | 0.21573988 |
| Treatment | 1 | 0.29537041 | 0.29537041 | 14.8758991 | 0.00014221 |
| OD:Genotype | 2 | 0.00438 | 0.00219 | 0.11029611 | 0.89560739 |
| OD:Treatment | 3 | 0.20477958 | 0.06825986 | 3.43780821 | 0.01735099 |
| Genotype:Treatment | 8 | 0.35585268 | 0.04448159 | 2.24025005 | 0.02477724 |
| OD:Genotype:Treatment | 6 | 0.18247745 | 0.03041291 | 1.53170172 | 0.167609 |
| Residuals | 283 | 5.61914442 | 0.01985563 | NA | NA |

F. **Fraction symptomatic on day 4**

|  | Df | Sum Sq | Mean Sq | F value | Pr(>F) |
| --- | --- | --- | --- | --- | --- |
| OD | 1 | 0.20280179 | 0.20280179 | 14.4481334 | 0.00017645 |
| Genotype | 2 | 0.06378955 | 0.03189477 | 2.27226763 | 0.10495579 |
| Treatment | 1 | 0.8422642 | 0.8422642 | 60.0051206 | 1.70E-13 |
| OD:Genotype | 2 | 0.02006727 | 0.01003363 | 0.71482253 | 0.49016019 |
| OD:Treatment | 3 | 0.09583971 | 0.03194657 | 2.2759578 | 0.08000022 |
| Genotype:Treatment | 8 | 0.22844396 | 0.0285555 | 2.03436866 | 0.04251409 |
| OD:Genotype:Treatment | 6 | 0.12316731 | 0.02052789 | 1.46246059 | 0.19102883 |
| Residuals | 283 | 3.97234047 | 0.01403654 | NA | NA |

**Table S4: Correlation matrix for different measures of pathogenicity containing R values corresponding to Figure 4B.**

|  | Fs_Day4 | PP_Day4 | Prop_Day4_Alive | Risk_Score | Fs_T | PP_T | Fs_T/T |
| --- | --- | --- | --- | --- | --- | --- | --- |
| Fs_Day4 | 1 | 0.43073123 | -0.7958791 | 0.66298673 | 0.68702919 | 0.29989318 | 0.69204839 |
| PP_Day4 | 0.43073123 | 1 | -0.490269 | 0.33487656 | 0.28629775 | 0.84079883 | 0.42863438 |
| Prop_Day4_Alive | -0.7958791 | -0.490269 | 1 | -0.8199348 | -0.5551074 | -0.3393062 | -0.8731669 |
| Risk_Score | 0.66298673 | 0.33487656 | -0.8199348 | 1 | 0.59044753 | 0.29626214 | 0.8503255 |
| Fs_T | 0.68702919 | 0.28629775 | -0.5551074 | 0.59044753 | 1 | 0.41849609 | 0.57822984 |
| PP_T | 0.29989318 | 0.84079883 | -0.3393062 | 0.29626214 | 0.41849609 | 1 | 0.30986907 |
| Fs_T/T | 0.69204839 | 0.42863438 | -0.8731669 | 0.8503255 | 0.57822984 | 0.30986907 | 1 |
